# Caecilian genomes reveal the molecular basis of adaptation, and convergent evolution of limblessness in snakes and caecilians

**DOI:** 10.1101/2022.02.23.481419

**Authors:** Vladimir Ovchinnikov, Marcela Uliano-Silva, Mark Wilkinson, Jonathan Wood, Michelle Smith, Karen Oliver, Ying Sims, James Torrance, Alexander Suh, Shane A. McCarthy, Richard Durbin, Mary J. O’Connell

**Affiliations:** Computational and Molecular Evolutionary Biology Group, School of Life Sciences, Faculty of Medicine and Health Science, University of Nottingham, NG7 2RD, UK; Tree of Life Programme, Wellcome Sanger Institute, Wellcome Genome Campus, Cambridge CB10 1SA, UK; Herpetology Laboratory, The Natural History Museum, London SW7 5BD, UK; Scientific Operations, Wellcome Sanger Institute, Wellcome Genome Campus, Cambridge CB10 1SA, UK; School of Biological Sciences, University of East Anglia, NR4 7TU, Norwich, UK; Department of Organismal Biology, Science for Life Laboratory, Uppsala University, SE-752 36, Uppsala, Sweden; Department of Genetics, University of Cambridge, Cambridge CB2 3EH, UK

**Author notes:** **Corresponding authors:** Mary O’Connell < >, Richard Durbin < >. both authors contributed equally (Vladimir Ovchinnikov, Marcela Uliano-Silva).

**Keywords:** Gymnophiona, Amphibia, Vertebrate Comparative Genomics, Limblessness

## Abstract

We present genome sequences for the caecilians *Geotrypetes seraphini* (3.8Gb) and *Microcaecilia unicolor* (4.7Gb), representatives of a limbless, mostly soil-dwelling amphibian clade with reduced eyes, and unique putatively chemosensory tentacles. More than 69% of both genomes are composed of repeats, with retrotransposons the most abundant. We identify 1,150 orthogroups which are unique to caecilians and enriched for functions in olfaction and detection of chemical signals. There are 379 orthogroups with signatures of positive selection on caecilian lineages with roles in organ development and morphogenesis, sensory perception and immunity amongst others. We discover that caecilian genomes are missing the ZRS enhancer of Sonic Hedgehog which is also mutated in snakes. *In vivo* deletions have shown ZRS is required for limb development in mice, thus revealing a shared molecular target implicated in the independent evolution of limblessness in snakes and caecilians.

---

Living amphibians, frogs, salamanders and caecilians, have diverged since the Triassic. They, or their ancestors, survived all mass extinctions including the Permian-Triassic which obliterated most terrestrial vertebrates (Wake and Vredenburg 2008). Our current extinction crisis places amphibians amongst the most threatened of the vertebrate groups (Blaustein and Wake 1990). In addition, the large and highly repetitive genomes representative of amphibia pose some of the greatest challenges for vertebrate genomics (Funk et al. 2018; Nowoshilow et al. 2018). Undoubtedly, reference quality genomes for amphibia will be important in addressing key aspects of their conservation, disease ecology and evolution, and breeding programs.

Caecilians (Gymnophiona) are the deepest diverging of the three extant amphibian orders and the sister group of the frogs and salamanders (Batrachia), diverging perhaps more than 300 MYA (Siu-Ting et al. 2019). Compared to batrachians, caecilians are few in number (approximately 215 species). With mostly secretive burrowing lifestyles and restricted distributions in the wet tropics west of Wallace’s line, they are relatively seldom encountered and often considered to be the least well-known group of tetrapods (Wilkinson 2012).

Caecilians are highly distinctive in their elongate (from 10 cm to 2 m adult lengths), and externally segmented snake- or worm-like form. Living species lack any trace of limbs or girdles, have skulls that are comparatively heavily ossified compared to batrachians, and have very short tails or no tails at all, all features that are associated with the fossorial or burrowing habits of adults. Eyes are also greatly reduced with any loss of vision seemingly compensated for by a putative chemosensory pair of tentacles on the snout that are not found in any other taxa (Wilkinson 2012, Taylor EH 1968). Other unique features include a dual-jaw closing mechanism, a copulatory organ formed from the hind part of the gut (phallodeum), and persistent Mullerian ducts in males. Their scientific name Gymnophiona means ‘naked snakes’ reflecting their perceived affinity to snakes albeit without scales. Ironically, some caecilians do have subdermal scales (quite different to the external scales of squamates) concealed in pockets or folds in the skin and are the only living amphibians to have scales. Like most other amphibians, caecilians are generalist predators as adults (Wilkinson 2012, Measey et al. 2004).

As with other living amphibians, oviparity with an aquatic larval stage and metamorphosis to a terrestrial adult is the ancestral reproductive mode within the group. Clutches of relatively few eggs are laid on land rather than in water, entailing a migration to water for any hatchling-larvae, and are invariably guarded until hatching by attending mothers. Other reproductive strategies include oviparity with direct terrestrial development and viviparity. Foetuses of at least some viviparous caecilians are believed to use specialised teeth to feed on the hypertrophied and lipidified oviduct linings of their mothers and it was discovered that in some oviparous direct developers their hatchlings feed on the similarly modified maternal epidermis with similarly specialised vernal teeth (Kupfer et al. 2006). Caecilian diversity is far from completely known and most of the described species are data deficient in the IUCN red list and thus lack any assessment of their conservation status and threats. New higher taxa (families and genera) have been recently discovered and caecilian species are described every year (Kamei et al. 2012; Wilkinson et al. 2021). Although many aspects of caecilian biology remain to be adequately investigated, phylogenetic relationships of the 10 currently recognised families are reasonably well-established, and support the generally accepted idea that caecilians are an ancient Gondwanan group with relatively recent and limited dispersals into Central America and South East Asia (Gower et al. 2002; Kamei et al. 2012).

The Rhinatrematidae, the deepest diverging (c. 125-MYA) of the ten caecilian families (Wilkinson et al. 2011), is represented by the only previously published caecilian genome *Rhinatremata bivittatum*, which is 5.3 Gb in size and was sequenced by the Vertebrate Genomes Project (VGP) (Rhie et al. 2021). Here we provide the reference quality genomes for two additional caecilian genomes, *Geotrypetes seraphini* (3.8 Gb) and *Microcaecilia unicolor* (4.7 Gb) and describe molecular level insights gleaned from their comparison with other vertebrate genomes.

## Reference genomes

The reference genomes of *Geotrypetes seraphini* (Dermopdiidae) and *Microcaecilia unicolor* (Siphonopidae) were assembled using 4 data types including Pacbio CLR and Hi-C reads, 10X Chromium linked-reads and BioNano optical maps (**Supplementary Table S1**) and meet the VGP’s 6.7.P5.Q40.C90 metric standards, the same used previously for *Rhinatrema bivittatum* and other vertebrates (Rhie et al. 2021). *G. seraphini* and *M. unicolor* respectively presented: contig N50 of 20.6 Mb and 3.6 Mb; scaffold N50 of 272 Mb and 376 Mb; and, Phred-scaled base accuracy Q43 and Q37 with 99% and 97% of sequences assigned to 19 and 14 chromosomes (**Table 1**). Chromosomal units were identified and named by size. The final assembly sizes were 3.8 Gb and 4.7 Gb, respectively **(Table 1**). Manual curation was performed as in Howe et al.(Howe et al. 2021) (**Supplementary Figure S1**) resulting in 69 and 55 removals of misjoins, 122 and 84 new joins, and 18 and 0 removals of false duplications for G. seraphini and M. unicolor respectively.

**Table 1:**
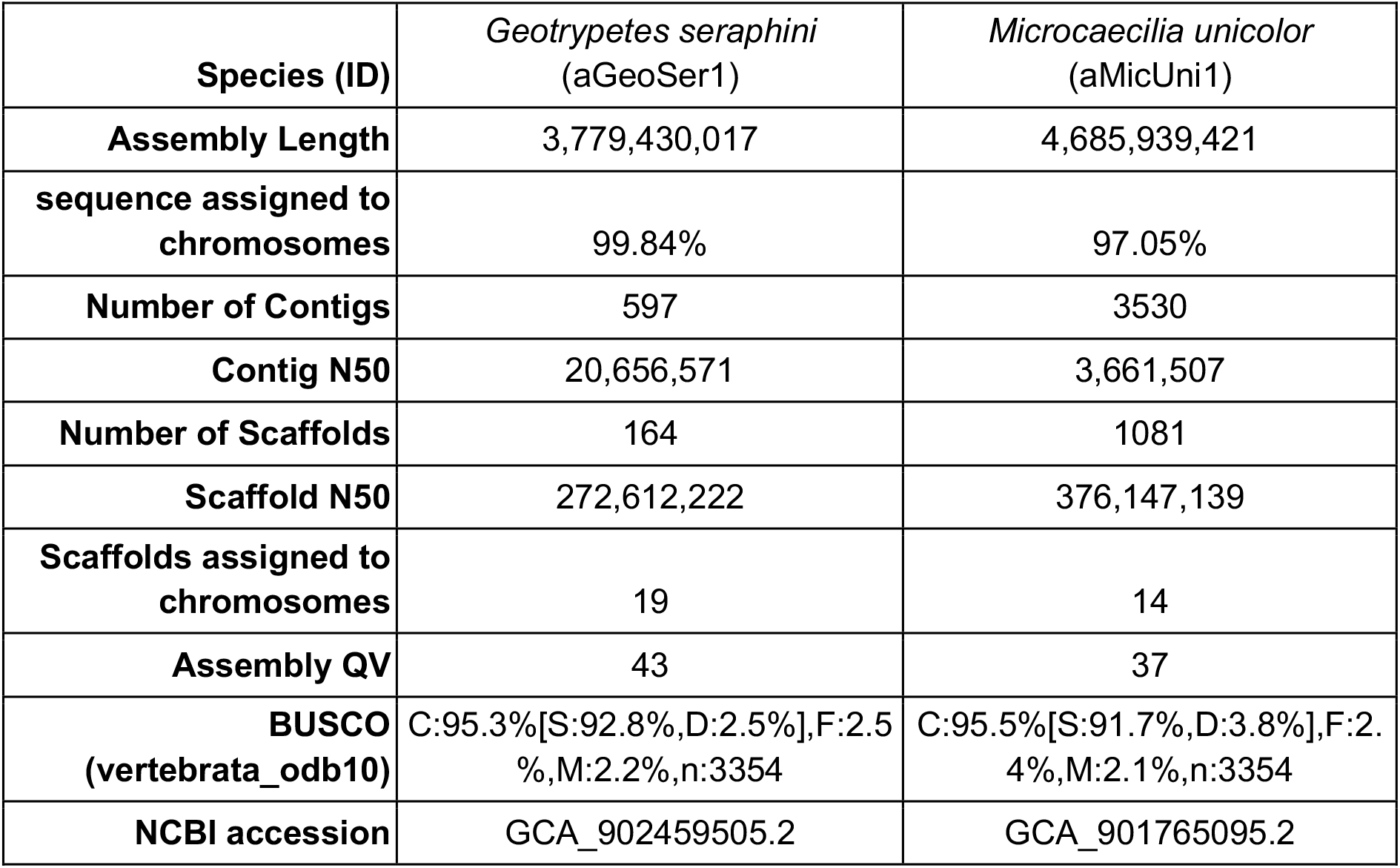
Final genome assembly statistics for *G. seraphini* and *M. unicolor*.

A synteny analyses performed with single-copy BUSCO genes shows that chromosome content and gene order are conserved to a remarkable extent across caecilian chromosomes, with large blocks of collinear synteny up to chromosome scale further conserved to anurans (common frog and toad) across more than 600 million years of evolution (**Figure 2**).

**Figure 1:**
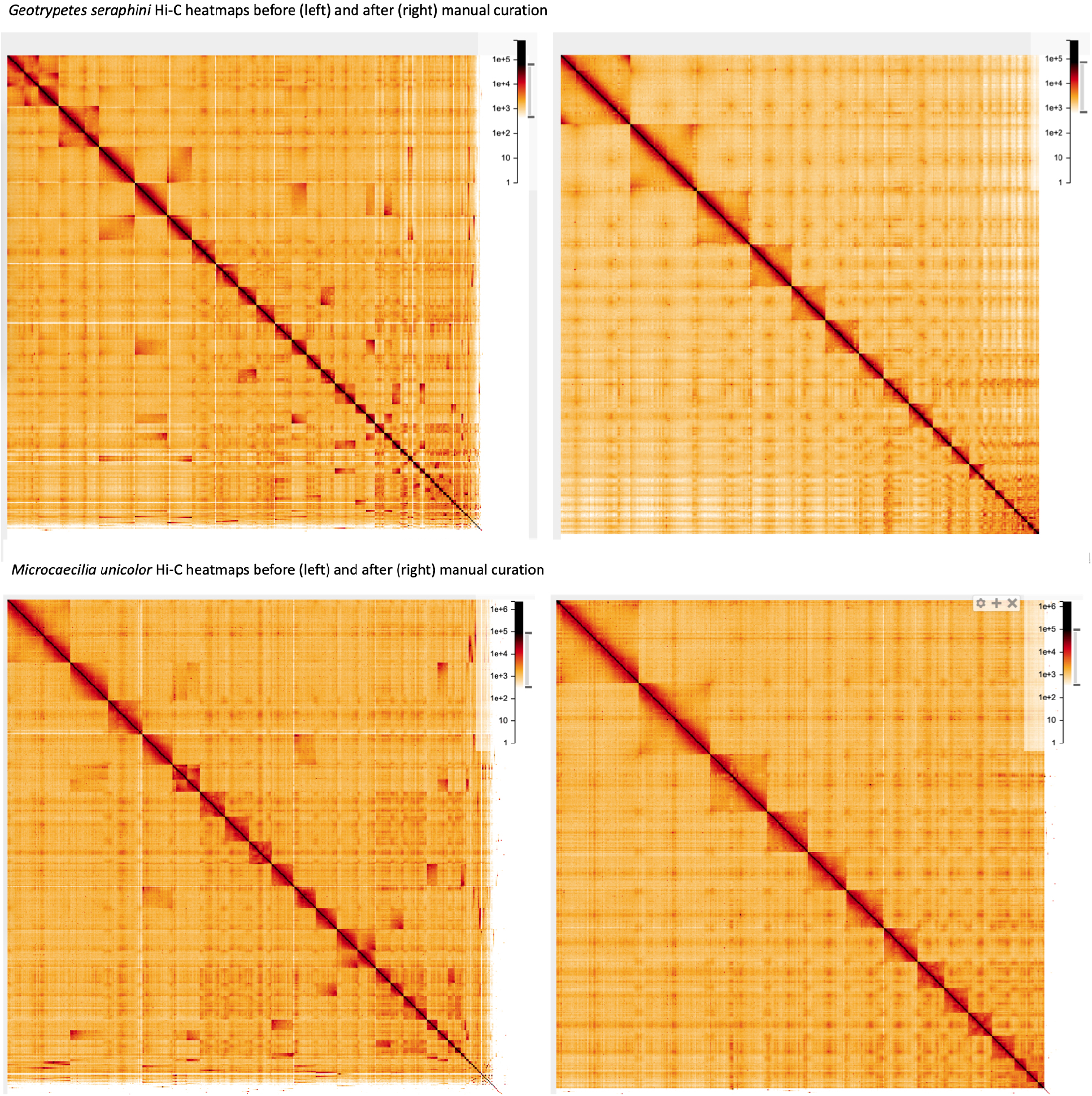
*G. seraphini* and *M. unicolor* genome HiC contact maps, respectively. The contact maps show Hi-C reads at 8.192 Mb resolution in HiGlass. The top two panels are *G. seraphini* and the bottom two panels are *M. unicolor* and, in both cases, the left panel is before and the right panel is after manual curation. Chromosomes are ordered from large (left/top) to small (right/bottom). After the VGP Assembly Pipeline and manual curation, 99.8% and 97% of sequences were assigned to 19 and 14 chromosomes for *G. seraphini* and *M. unicolor* respectively.

**Figure 2:**
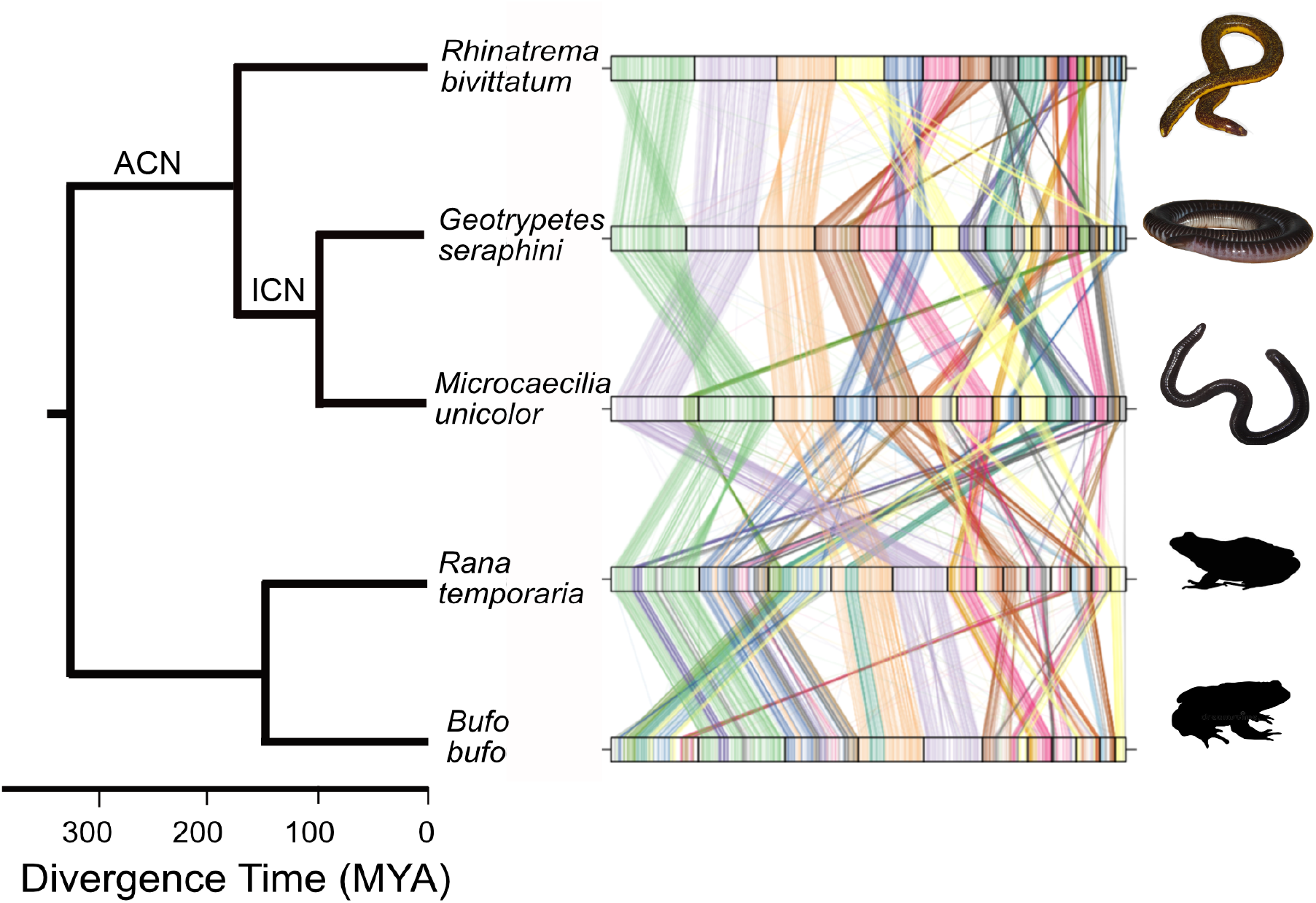
Synteny plots showing the conservation of large scale gene linkage and gene order across caecilians, and to a substantial extent across amphibia. Conserved unique single copy vertebrate genes were identified with BUSCO and connected by lines coloured according to their chromosomal location in *Rhinatrema bivittatum*. The Ancestral Caecilian Node (ACN) and Internal Caecilian Node (ICN) are labelled. Common frog *Rana temporaria* and toad *Bufo bufo* genomes from https://wellcomeopenresearch.org/articles/6-286 and https://wellcomeopenresearch.org/articles/6-281, respectively. Synteny was identified with ChrOrthLink (https://github.com/chulbioinfo/chrorthlink). Images of caecilians are modified using the Gimp software from original photos taken by Mark Wilkinson. Frog and toad silhouettes are taken from http://phylopic.org/.

## Repeat content

Substantial proportions of the caecilian genomes were found to consist of repeats: a total of 67.7%, 72.5% and 69.3% for *R. bivittatum* (Rhie et al. 2021), *G. seraphini* and *M. unicolor* respectively (**Supplementary Table S2**). Class I transposable elements (TEs; retrotransposons) are ∼20 times more abundant (in base pairs) than Class II TEs (DNA transposons) and make up more than 30% of each caecilian genome. Long INterspersed Elements (LINEs) are the most abundant transposon type, followed by Dictyostelium Intermediate Repeat Sequences (DIRSs), *i*.*e*. tyrosine recombinase retroelements. These relative proportions differ from those found in the large genomes of other amphibians including caecilians; for example, a genomic low-coverage shotgun analysis of the caecilian *Ichthyophis bannanicus* (genome size 12.2 Gb) revealed more DIRSs than LINEs (Wang et al. 2021), while published salamander genomes are dominated by Long Terminal Repeat (LTR) elements, with DIRSs never surpassing 7% of their content (Sun and Mueller 2014; Nowoshilow et al. 2018). These findings bolster the concept that repeated extreme TE accumulation in amphibians is not resulting from failure to control a specific type of TE (Wang et al. 2021).

## Gene family analyses

Comparing the protein coding regions of the three caecilian genomes across 22 vertebrate genomes we identified a set of 31,385 orthogroups, of which 15,216 contained caecilian genes. We identified 265 gene families present across vertebrates but missing in amphibia, and an additional 260 orthogroups lost specifically in caecilians (**Supplementary Table S3**). In contrast, 1,150 orthogroups are present only in caecilians (**Supplementary Table S4**) and are enriched for functions such as olfaction and detection of chemical signals (p-value<0.01). At least 20% of these caecilian specific genes contained one of three protein domains (zf-C2H2, KRAB, 7tm_4). The 7tm_4 proteins are transmembrane olfactory receptors (Buck and Axel 1991); enrichment of this domain amongst the novel protein families in caecilians suggests an intense selective pressure on chemosensory perception at the origin of the caecilians, as they adapted to life underground with reduced vision and compensatory elaboration of chemosensory tentacles. Proteins containing zf-C2H2 and KRAB domains are known to have functions in regulating transcription, with zf-C2H2 containing proteins in humans shown to recognize more motifs than any of the other transcription factors combined. In addition, KRAB and zf-C2H2-containing proteins have been shown to bind currently active and ancient families of specific TEs (*e*.*g*. LINEs and LTRs/ERVs) (Najafabadi et al. 2015). The emergence of novel gene families with these functional capacities at the origin of caecilians may have contributed to the unique pattern of TE accumulation we observe in this group; further work is needed.

We performed a gene birth and death analysis using CAFE v5 (Mendes et al. 2020) on the remaining 13,541 orthogroups, examining the ancestral and extant caecilian nodes where possible. The majority of these (10,035) orthogroups had no net change in gene family size between caecilian species and the ancestral amphibian node (8,065 orthogroups) or had insufficient sampling (1,970 orthogroups), and were excluded from further analysis. We reconstructed ancestral states for the remaining 3,506 orthogroups (**Supplementary Table S5**). There were 156 orthogroups that were completely absent in *G. seraphini* and *M. unicolor* (most likely lost in their most recent common ancestor) (**Supplementary Table S3**). Only 13 orthogroups showed significant changes in number in caecilians (**Figure 3, Supplementary Table S6**), with 5 expansions at the ancestral caecilian node (ACN), and 3 at the internal caecilian node (ICN), of which one gene family is significantly expanded at both nodes. There are a total of three gene families with significant contractions, all of which are on the ACN. The gene families displaying significant expansions are: cytochrome P450 family 2 (ACN), these monooxygenases catalyse many reactions involved in metabolism of a large number of xenobiotics and endogenous compounds (Manikandan and Nagini 2018); butyrophilin (BTN) family (ACN), involved in milk lipid secretion in lactation and regulation of the immune response (Afrache et al. 2012); tripartite motif (TRIM) family (ACN and ICN) involved in a broad range of biological processes that are associated with innate immunity (Ozato et al. 2008); and H2A and H2B histones (ICN), which together with H3 and H4 histones and DNA form a nucleosome (Koyama and Kurumizaka 2018). In contrast, while immune function related butyrophilin and TRIM families have significant expansions at the ACN and/or ICN, both immunoglobulin heavy and light variable gene families have significant contractions at the ACN. The final gene family displaying significant contractions is gamma crystallin, a structural protein found largely in the nuclear region of the lens of the eye at very high concentrations (Vendra et al. 2016). Changes in these gene family repertoires may have contributed to the transition to a fossorial lifestyle and packaging of a large genome.

**Figure 3:**
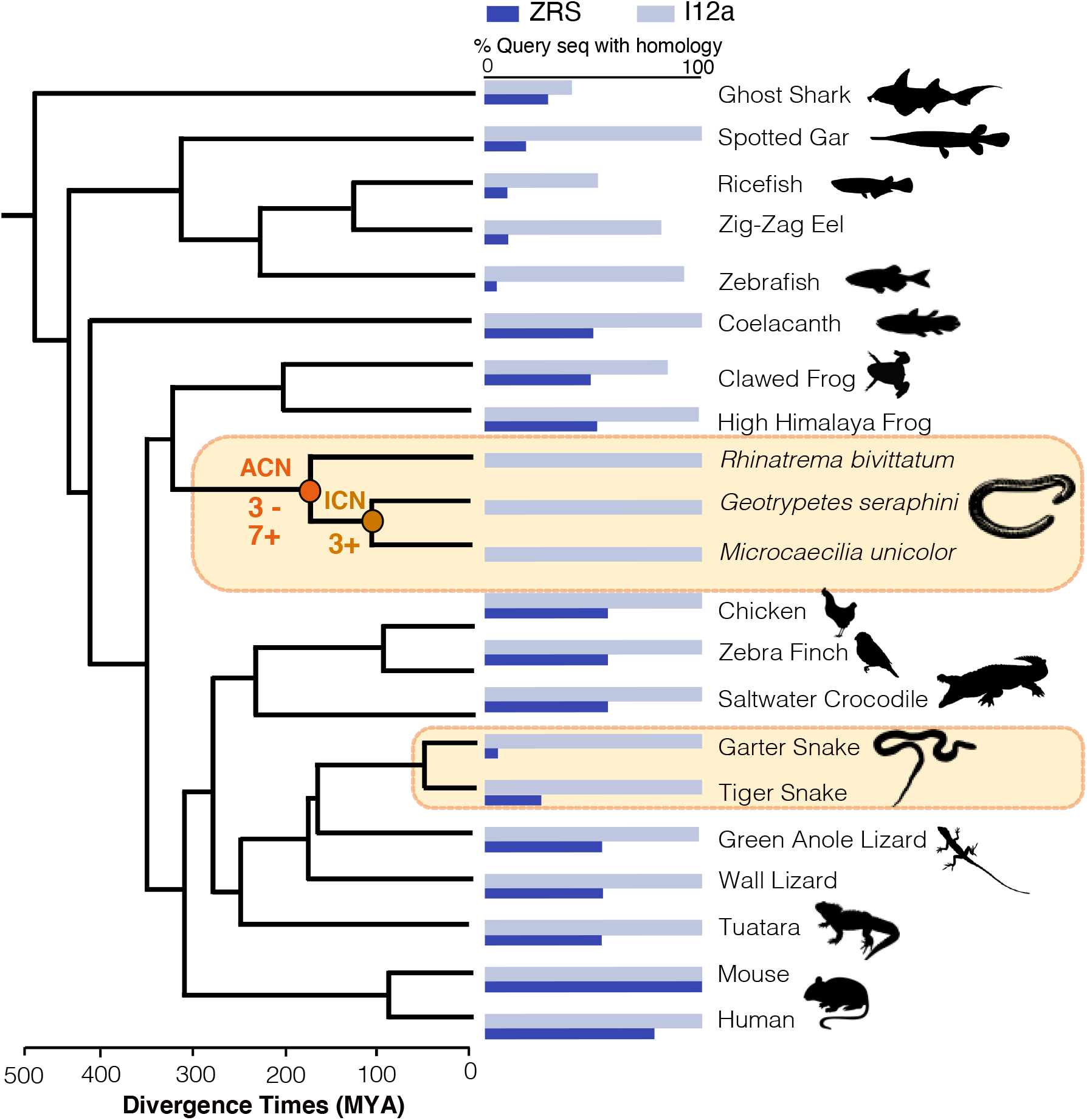
Summary of sequence conservation of two enhancer elements across vertebrates (ZRS and l12a). The vertebrate species phylogeny used throughout this study is shown on the left with the significant gene gain and loss events noted on the ancestral and internal caecilian nodes (ACN and ICN), respectively. The histogram shows the level of sequence conservation identified by BLASTN for each species for two enhancers: I12a (light blue) and ZRS (dark blue). Snakes and caecilians are highlighted as they independently evolved limbless morphologies. Animal images are taken from http://phylopic.org/.

## Identification of genes with signatures of positive selection

Variation in selective pressure was assessed using codon-based models of evolution to assess changes in dN/dS across sites and lineages as implemented in codeml in the PAML package (Yang 2007). All 1,935 gene families that reached our criteria (see materials and methods) were analysed. These 1,935 gene families were functionally enriched for extracellular structure organization, developmental process, regulation of biological process, response to stress, cell communication, signal transduction, regulation of signalling, and leukocyte differentiation (**Supplementary Table S7**). The lineages specified as foreground were the branch leading to the extant caecilians (ACN), and all terminal and internal branches within the caecilian clade. The selective pressures, *i*.*e*. positive or negative selection or neutral evolution, were estimated for each gene within each foreground lineage and were compared to all other vertebrates (background lineages) in the alignment. Here we report the signatures of positive selection (dN/dS >1) identified in homologs on the foreground (*i*.*e*. caecilian) lineages. After Bonferroni correction, we detected 379 orthologous families with evidence of caecilian lineage-specific positive selective pressure (**Supplementary Table S8**). We did not find any statistical enrichment for GO functions in the genes under positive selection on the nodes tested. Examples of genes with signatures of positive selection on the ACN are: FBN1 (under positive selection on both the ACN and the ICN), AGTPBP1, and CEP290 all of which are involved in eye morphogenesis (Chakrabarti et al. 2006; Sheck et al. 2018; Stephenson et al. 2020). Genes with signatures of positive selection on the ICN include: THBD (involved in the reduction of thrombin), WLS (enables Wnt-protein binding activity and is involved in several processes, including animal organ development; mesoderm formation; and positive regulation of canonical Wnt signalling pathway), and CD8A (mediates efficient cell-cell interactions within the immune system). In addition, COL3A1 is under positive selection on all caecilian nodes tested (*i*.*e*. ACN, ICN and terminal nodes). A sample of the genes under positive selection within specific caecilian lineages are described as follows (specific internal caecilian lineage in parenthesis): HESX1 (*R. bivittatum*) required for the normal development of the forebrain, eyes and other anterior structures such as the olfactory placodes and pituitary gland (Dattani et al. 1998: 1); NFE2L2 (*G. seraphini*), a transcription factor that plays a key role in the response to oxidative stress (Huang et al. 2000; Eggler et al. 2009; Huppke et al. 2017; Sanghvi et al. 2019); LGR4 (*R. bivittatum*) is involved in the development of the anterior segment of the eye (Siwko et al. 2013) and is required for the development of various organs, including kidney, intestine, skin and reproductive tract (Hoshii et al. 2007; Kinzel et al. 2014); COL9A3 (*M. unicolor*) encodes a component of Collagen IX - a structural component of cartilage, intervertebral discs and the vitreous body of the eye (Olsen 1997; He and Karsdal 2016). In summary, whilst the biological processes and functions of the genes under positive selection are not significantly enriched, there are several genes implicated in organ (especially eye) development and morphogenesis. Caecilian tentacles can be considered as compensation for reduced vision through enhanced olfaction, and they are thought to be materially related (by transformation) to components of the visual system such as eyelids and lacrimal ducts (Billo and Wake 1987). Therefore, a tentative explanation for the positive selection we observe on genes associated with organ (eye) development and morphogenesis is the origin of the tentacles from ancestrally visual components. The approach we have taken in our analysis of selective pressure variation is necessarily stringent, and therefore not a complete assessment of the entire genome where there are likely many other processes at work.

## Analysis of ZRS enhancer loss

Some key enhancers for developmental regulator genes are very strongly conserved at the sequence level across all vertebrates. For example, the I12a enhancer element, located between homeobox genes Dlx1 and Dlx2, is known to be conserved from bony fish to mice (Plessy et al. 2005). Analysis of the ortholog of the l12a enhancer across the 22 vertebrate species confirms that it is easily identifiable and conserved in all vertebrates, including the three caecilians (**Figure 3**). Similarly, the ZRS enhancer element for the Sonic hedgehog gene (Shh), which is located within an intron of the LMBR1 gene, is almost ubiquitously conserved in vertebrates. However, snakes contain a mutant form of ZRS that when placed into mice produces a “serpentised” phenotype, directly implicating loss of ZRS function in vertebrate limblessness (Kvon et al. 2016). From the fossil record we know that snake limblessness pre-dates that of limbless lizards, also reflected in a higher level of divergence in limb regulatory elements in snakes in comparison to limbless lizards. Indeed, ZRS is intact in limbless lizards where more complex and lineage-specific routes to limblessness have been proposed (Roscito et al. 2022). Here we show that the conserved ZRS element is absent (or mutated beyond recognition) in the three caecilian genomes. Specifically, there is no trace of homology by sequence matching (**Figure 3**), and a conserved ETS1 binding site within the ZRS enhancer element, which has been shown to be critical for limb development in mouse and is missing in snakes (Kvon et al. 2016), is also entirely missing in caecilians (**Supplementary Figure S2**). Combined with the functional work on the mutated form of the snake ZRS, this may provide us with a potential common molecular target implicated in the convergent loss of limbs in snakes and caecilians. Alternatively, the observed pattern of loss of the ZRS element in caecilians could be secondary to the loss of limbs. Similar to the situation in lizards (Roscito et al. 2022), loss of limbs in caecilians could have been piecewise and relaxation of selective pressures on the ZRS region could have resulted in its eventual loss from caecilian genomes. Functional analysis will be needed to finally resolve the history of limb loss in this major amphibian group.

## Materials and Methods

### Sample preparation and genome assembly

Genome sequences were produced from wild-caught animals that had been maintained in captivity for several years. Specimens are at the Natural History Museum, London catalogued under their unique field tags: *G. seraphini* (MW11051) from Kon, Cameroon, and *R. bivittatum* (MW11052) and *M. unicolor* (MW11053), both from Camp Patawa, Kaw Mountains, French Guiana. All DNA extractions were from liver tissue using the Bionano Animal Tissue Plug preparation (https://bionanogenomics.com/wp-content/uploads/2018/02/30077-Bionano-Prep-Animal-Tissue-DNA-Isolation-Soft-Tissue-Protocol.pdf). Pacific Biosciences libraries were prepared with the Express Template Prep Kit 1.0 and Blue Pippin size selected. Pacific Biosciences CLR data was generated from 36 SMRTcells of *M. unicolor* and 6 SMRTcells of *G. seraphini* sequenced with the S/P2-C2/5.0 sequencing chemistry on the Pacific Biosciences Sequel machine. A further 5 SMRTcells of *G. seraphini* were sequenced with S/P3-C1/5.0-8M sequencing chemistry on a Pacific Biosciences Sequel II machine. The Hi-C libraries were created with a Dovetail Hi-C kit for *G. seraphini* and an Arima Genomics kit (version 1) for *M. unicolor* and sequenced on an Illumina HiSeq X. A 10X Genomics Chromium machine was used to create the linked-read libraries which were sequenced on an Illumina HiSeq X. Optical maps were created for both species using a Bionano Saphyr instrument.). Raw reads statistics and data access links are available in **Supplementary Table S1**.

Assembly for *G. seraphini* and *M. unicolor* was conducted mainly as for *R. bivittatum* as described in (Rhie et al. 2021) using four data types and the Vertebrate Genomes Project (VGP) assembly pipeline (version 1.6 for *G. seraphini* and version 1.5 for *M. unicolor*; **Supplementary Figure S1**). In brief, the Pacific Biosciences CLR data for each species was input to the diploid-aware long-read assembler FALCON and its haplotype-resolving tool FALCON-UNZIP (Chin et al. 2016). The resulting primary and alternate assemblies of *M. unicolor* were input to Purge Haplotigs (Roach et al. 2018) and *G. seraphini* assemblies were input to Purge_dups (Guan et al. 2020) for identification and removal of remaining haplotigs. Both species’ primary assemblies were subject to two rounds of scaffolding using 10X long molecule linked-reads and Scaff10X (https://github.com/wtsi-hpag/Scaff10X), and one round of Bionano Hybrid-scaffolding with pre-assembled Cmaps from 1-enzyme non-nicking (DLE-1) and the Solve Pipeline. The resulting scaffolds were then further scaffolded into chromosome-scale scaffolds using the Dovetail/Arima library Hi-C data for *G. seraphini/M. unicolor* and SALSA2 (Ghurye et al. 2019). The scaffolded primary assemblies plus the Falcon-phased haplotigs were then subjected to Arrow (Chin et al. 2013) polishing with the Pacbio reads and two rounds of short read polishing using the 10X Chromium linked reads, longranger align (Bishara et al. 2015), freebayes (Garrison and Marth 2012) and consensus calling with bcftools (Danecek et al. 2021) (further details available in Rhie et al 2021, and **Supplementary Figure S1**). Assemblies were checked for contamination and were manually curated using gEVAL system (Chow et al. 2016), HiGlass (Kerpedjiev et al. 2018) and PretextView (https://github.com/wtsi-hpag/PretextView) as described previously (Howe et al. 2021). Mitochondria were assembled using mitoVGP (Formenti et al. 2021). Manual curation was performed as described in Howe et al. (Howe et al. 2021). Genome annotation was carried out using the NCBI Eukaryotic Genome Annotation Pipeline, which produces homology-based and *ab initio* gene predictions to annotate genes (including protein-coding and non-coding as lncRNAs, snRNAs), pseudo-genes, transcripts, and proteins (https://www.ncbi.nlm.nih.gov/genbank/eukaryotic_genome_submission_annotation/). Caecilian annotations available on NCBI at the accessions GCF_902459505.1, GCF_901765095.1 are summarised in **Supplementary Table S9**. Raw reads statistics, accession numbers and software versions employed can be found at **Supplementary Table S1**.

Prediction and annotation of repeats was achieved using a *de novo* library of repeats generated with RepeatModeler2 for each species (Flynn et al. 2020), combined with the Repbase “Amphibia” library (release 26.04) (Bao et al. 2015) to form the final library for each species. Repeats were masked with RepeatMasker (http://www.repeatmasker.org/) and Window Masker (Morgulis et al. 2006). Then transcripts, proteins and RNA-Seq from the NCBI database were aligned to the genomes using Splign (Kapustin et al. 2008) and ProSplign (https://www.ncbi.nlm.nih.gov/sutils/static/prosplign/prosplign.html). Alignments were submitted to Gnomon (https://www.ncbi.nlm.nih.gov/genome/annotation_euk/gnomon/) for gene prediction. Models built on RefSeq transcript alignments were given preference over overlapping Gnomon models with the same splice pattern. **Supplementary Table S2** presents a summary of caecilian repeat annotations. RepeatModeler libraries in fastA format are available from DOI:10.5281/zenodo.7540729.

### Data assembly and treatment for the comparative study

Coding DNA sequences (CDSs) for 21 vertebrate species (**Supplementary Figure S3**) were downloaded from Ensembl release 100 (Yates et al. 2020). In those cases where a more contemporary version of the genome was available on RefSeq (Release 200) (O’Leary et al. 2016) we used the RefSeq genome and corresponding annotations (**Supplementary Table S10**). The longest canonical protein coding region for each gene was retained for further analysis.

### Orthogroup prediction and gene birth and death analysis

We identified 31,385 orthogroups for the 419,877 protein coding regions across 21 vertebrate species using OrthoFinder (Emms and Kelly 2019) (all orthogroups are available at DOI:10.5281/zenodo.7540729). We used a phylostratigraphic approach to explore caecilian specific losses in the context of the uncontroversial vertebrate phylogeny used throughout (**Figure 3**), which we extracted from timetree.org (Kumar et al. 2017) assembled from the following literature: caecilians (San Mauro et al. 2014), amphibians (Siu-Ting et al. 2019), fish (Betancur-R et al. 2017), reptiles (Pyron et al. 2013), mammals (Morgan et al. 2013; Tarver et al. 2016), and birds (Chiari et al. 2012). The phylogenetic distribution of the orthogroups revealed 1,150 that were gained in caecilians, and 525 that were absent in all three caecilians. Information about species-specific losses elsewhere in the tree was not carried forward for further analysis. We partitioned the orthogroups that lack caecilian representation in the following ways: (1) to identify orthogroups that lack representation across all amphibia: we identified orthogroups that contained at least two fish species and two tetrapod (non-amphibian) species - totalling 265 orthogroups, (2) to identify orthogroups that are absent only in caecilians: we extracted those orthogroups with least two fish species and two tetrapod species (including at least one frog species) - totalling 238 orthogroups, (3) to identify orthogroups that are present across amphibia and amniota but absent in caecilians: we extracted orthogroups containing two frog species and two amniota species - totalling 22 orthogroups. Orthogroups that did not satisfy these filters were set aside. Combining the set of orthogroups that contain caecilian representatives (13,541) plus those that passed our filters 1-3 above (525), produced our final set of 14,066 orthogroups for analysis in CAFE v5 with Poisson distribution option and the lambda parameter (rate of change of evolution) estimated for each species (Mendes et al. 2020). All 3,506 orthogroups showing expansions or contractions within caecilians are provided in **Supplementary Table S5**, and orthogroups with significant expansions and contractions are detailed in **Supplementary Table S6**.

### Analysis of selective pressure variation

Our selective pressure variation analysis focussed on 3,236 single-copy orthogroups (SGOs) and 9,690 multi-copy genes (MCGs) from our orthogroups. The 9,690 MCGs obtained from the CAFE analysis, could be further broken down into SGO clusters as follows: 3,464 contained species specific duplications in a single lineage, and were designated SGOs by removal of the single lineage containing the duplications; the remaining 6,226 were divided into their constituent single-copy paralogous groups using UPhO (Ballesteros and Hormiga 2016). Species-specific gene duplications that were not specific to caecilians were removed. In total this provided 14,807 SGOs (3,236 original SGOs plus 11,571 SGOs generated from MCGs) for further analysis. We used three different alignment methods on the amino acid sequences for these SGOs (*i*.*e*. MAFFT (Rozewicki et al. 2019) (with –auto option), MUSCLE (Edgar 2004), and Prank (Löytynoja 2014) (with -nobppa option)), and used MetAl (Blackburne and Whelan 2012) to assess the statistical significance of the resultant alignments. If the difference between alignments was >=5%, the alignment with the highest NorMD (Thompson et al. 2001) score was used. The corresponding gene trees were reconstructed using IQtree (Nguyen et al. 2015) (with 100 bootstraps for each tree and models of best fit selected on a gene-by-gene basis). Robinson-Foulds distances between each of the gene trees generated and the canonical species tree were estimated using Clann (Creevey and McInerney 2005), and only those gene trees with zero distance were retained for further analysis, *i*.*e*. the gene and species tree were required to be in full agreement thus minimising the risk of hidden paralogy in our single-copy gene orthogroups (SGOs). It has been shown that codeml provides more accurate predictions when a minimum of 7 species are present in the dataset (Anisimova et al. 2002), gene families that did not meet this criterion were not considered for selective pressure variation analysis. We assessed the patterns of selective pressure variation on the remaining 1,935 SGOs using codon-based models of evolution in codeml (Yang 2007) using our pipeline for large scale analyses “Vespasian” (Constantinides et al. 2021). The models we employed are a set of standard nested models which are automatically compared by Vespasian using likelihood ratio tests with significance calculated using the appropriate degrees of freedom. The models used were the neutral model M1Neutral, and its lineage-specific extensions model A, and the null model for model A. M1Neutral allows two site classes for dN/dS (referred to as ω throughout): ω0=0 and ω1=1. Model A assumes the two site classes are the same in both foreground and background lineages (ω0=0 and ω1=1) and ω2 for the foreground is estimated from the data and free to vary above 1. Model A null estimates a ω2 value also, but here it is restricted to below 1 thus allowing sites to be evolving under either purifying selection, or to be neutrally evolving but not permitting positive selection. Sequences were considered to exhibit lineage-specific selective pressure if the likelihood ratio test for ModelA is significant in comparison to both ModelA null and M1Neutral. All alignments (codon-based and amino-acid) for the selective pressure analyses are available at DOI:10.5281/zenodo.7540729. The GO terms were predicted for all caecilian CDSs using eggNOG with “orthology restrictions” option set to “transfer annotations from one-to-one orthology only” (eggnog-mapper.embl.de) (Huerta-Cepas et al. 2019) and all other parameters as default. GO term enrichment analysis was carried out using goatools (Klopfenstein et al. 2018) with Taxonomic Scope auto-adjusted per query.

### Comparative analysis of the ZRS enhancer

The ZRS enhancer sequence is located within an intron between exons 5 and 6 of the mouse LMBR1 gene sequence (Gene ID: 105804842) (Kvon et al. 2016). The LMBR1 sequence was extracted from the genomes of each species in our sample set (**Supplementary Table S11**) and the homologous intron sequence containing the ZRS sequence was identified across all species. Using BLASTn (Camacho et al. 2009) the ZRS region was readily identifiable across all 22 non-caecilian species (**Figure 3**, and **Supplementary Figure S2**) but was not detectable in the three caecilian genomes. The ZRS sequence was also searched against the reference genome assemblies of all three caecilians (to account for possible relocation of the enhancer) and we did not identify a ZRS-like sequence in an alternative location in the caecilian genomes. Using the same approach, we quantified the level of sequence conservation across our set of vertebrates for an additional enhancer, I12a (AF349438.2), located between the homeobox bigene cluster paralogs DLX1 and DLX2 (**Supplementary Table S11**). The DLX1 gene was not annotated for Crocodylus porosus, therefore we used the region between METAP1D and DLX2.

## Acknowledgements

MUS, YS, JW, JT, KO, MS are supported by Wellcome grant WT206194, SAM and RD are supported by Wellcome grant WT207492, MW thanks the Direction de l’Environment de l’Aménagement et du Logement and Le Comité Scientifique Régional du Patrimonie Naturel, French Guiana and many colleagues for help in obtaining specimens. MJO’C would like to thank the University of Nottingham for awarding funds to support this work. MJO’C and VO are grateful for access to the University of Nottingham’s Augusta HPC service. For the purpose of open access, and as this research was funded in part by the Wellcome Trust [Grant numbers WT206194 and WT207492], the author has applied a CC BY public copyright licence to any Author Accepted Manuscript version arising from this submission.

## Author contributions

MW supplied all biological samples and contributed to the interpretation of results. MS performed DNA extractions and optical mapping. KO coordinated the creation of sequencing libraries and genomic sequencing. SAM generated the genome assemblies. YS, JT and JW performed the manual curation of the assemblies. MUS performed BUSCO synteny analyses and with AS the repeat analyses. MJO’C and VO performed and interpreted the selective pressure analyses and birth and death analyses. MJO’C and VO carried out the comparative analysis of ZRS and l12a enhancer elements and interpreted results. RD supervised the genomics aspects of the project and MJO’C the comparative analyses. All authors contributed to writing the manuscript.

## Supplementary Figures and legends

**Supplementary Figure S1**: Standard Vertebrate Genome Project (VGP) assembly pipeline used to assemble *G. seraphini* and *M. unicolor* genomes (vs 1.1-1.6). This diagram is taken from Rhie et al 2021.

**Supplementary Figure S2:** Cartoon of a 26bp fragment of the alignment of ZRS enhancer region across a range of vertebrates. The dashed lines indicate regions where the homologous sequence is not detectable, showing independent loss in snakes and caecilians of an otherwise well conserved tetrapod ZRS region. The yellow box shows the location of a critical ETS1 binding site within the ZRS1 enhancer element. This ETS1 binding site has been suggested to directly activate the ZRS enhancer by binding to multiple ETS recognition sites (Lettice et al. 2012). The precise location of the 17bp region “snake-specific deletion” shown to be important in limb development by Kvon et al (Kvon et al. 2016) is also denoted.

## Supplementary Tables

**Supplementary Table S1:** Final genome assembly statistics for three caecilian genomes.

**Supplementary Table S2:** Repeat classes found in caecilian genomes.

**Supplementary Table S3**: Orthogroups with total losses of genes in amphibians or caecilians. **Supplementary Table S4:** Caecilian-specific orthogroups and their constituent gene members. **Supplementary Table S5**: Orthogroups with their associated increase or decrease in gene numbers within caecilians.

**Supplementary Table S6**: Number of orthogroups with significant changes in gene family content in caecilians.

**Supplementary Table S7:** GO terms enriched in the 1,935 genes analysed for selective pressure variation.

**Supplementary Table S8:** Orthologous families with evidence of positive selective pressure in caecilians.

**Supplementary Table S9:** Annotation features described using the NCBI Eukaryotic Genome Annotation Pipeline.

**Supplementary Table S10:** Source of data for each species sampled, and the number of coding DNA sequences (CDSs) available per species.

**Supplementary Table S11:** Sequence data used for comparison of the ZRS enhancer region across the 21 vertebrate species used.

## Notes

https://zenodo.org/record/5780326/export/json#.YhTCq_XP0dc

### Competing Interest Statement

The authors have declared no competing interest.

### Summary of Updates

We have substantially updated the manuscript text and figures.

