## Supplementary figures and images for "Caecilian genomes reveal the molecular basis of adaptation, and convergent evolution of limblessness in snakes and caecilians"

### Figure 2

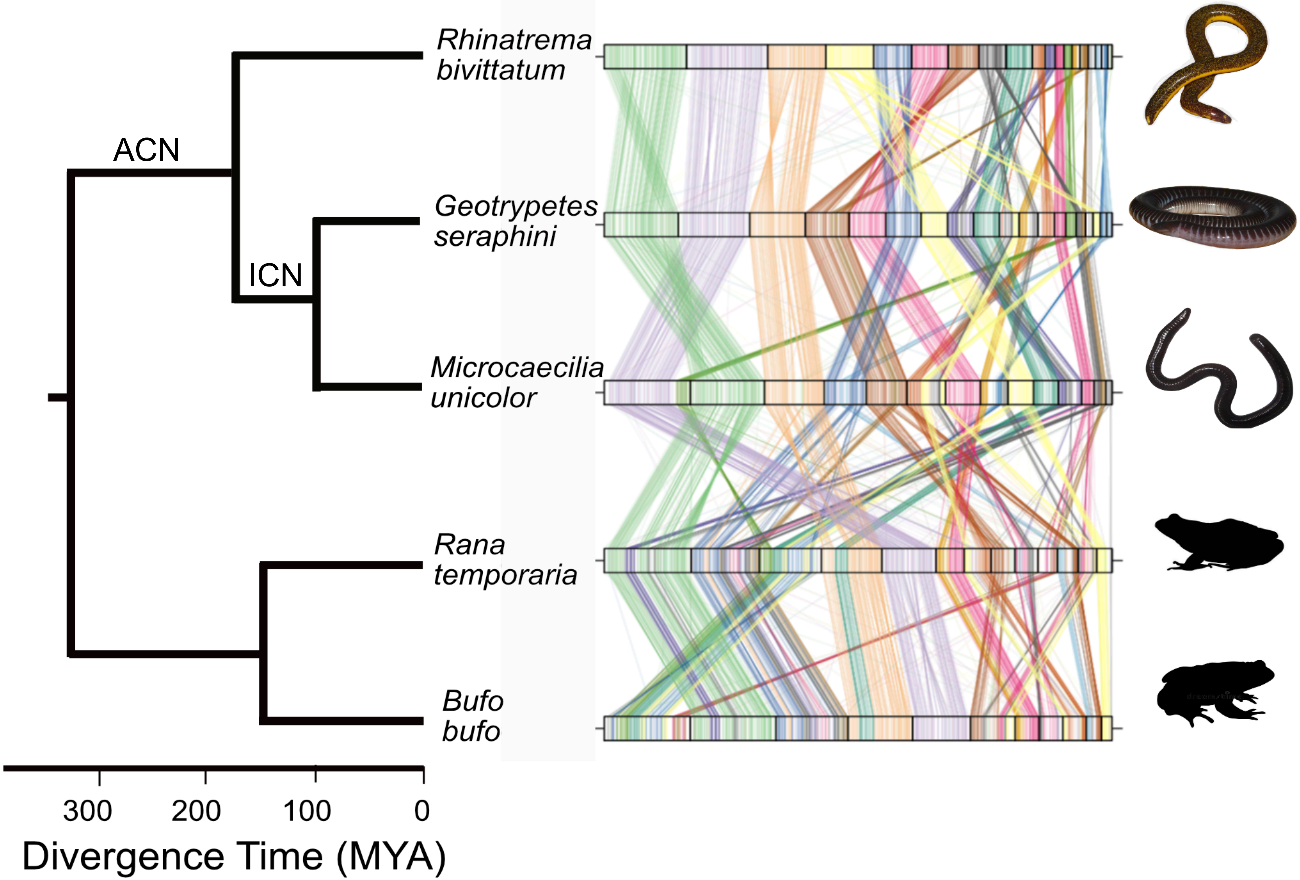

### Figure 3

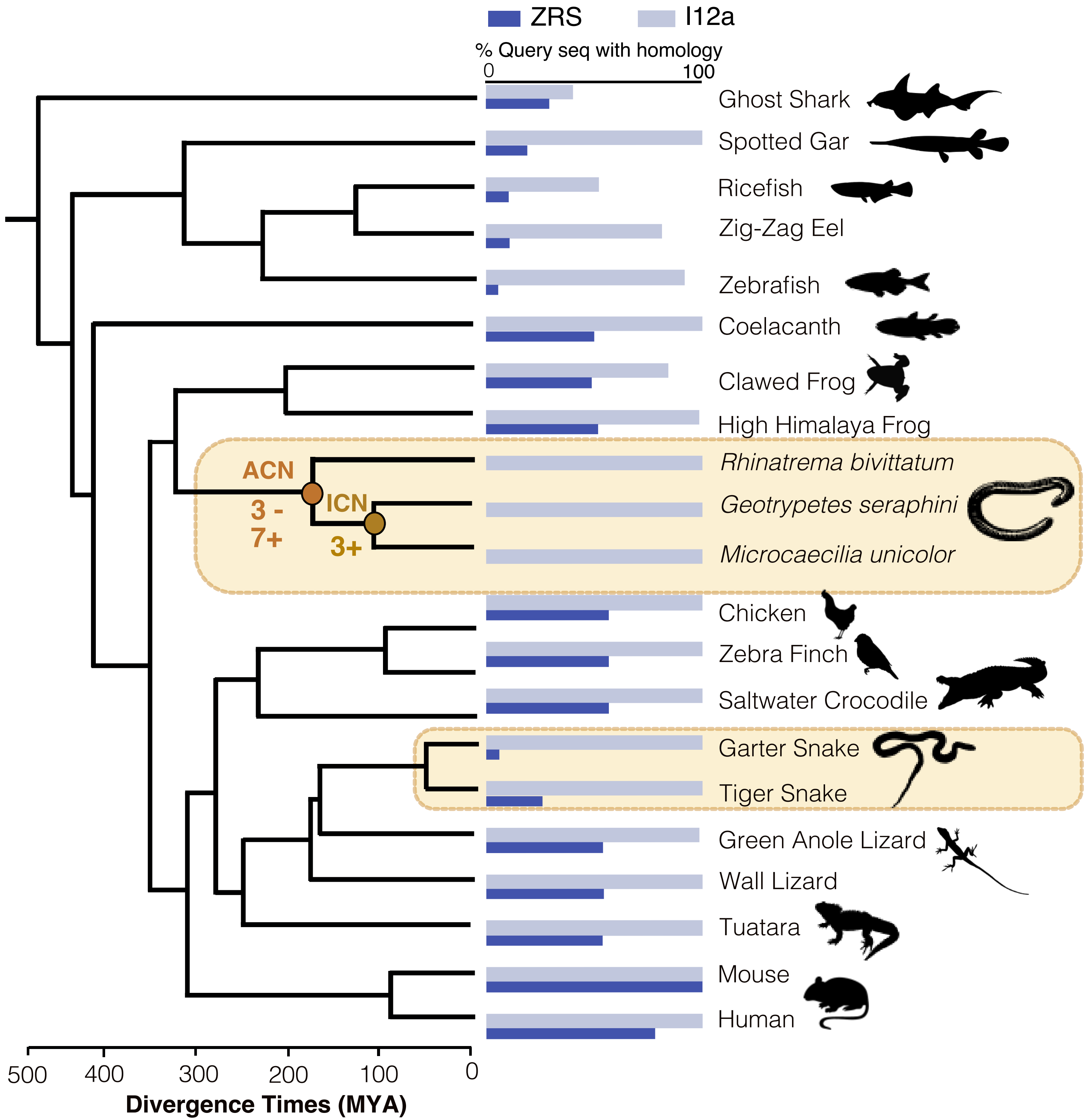

### supplementary Figure S1

a

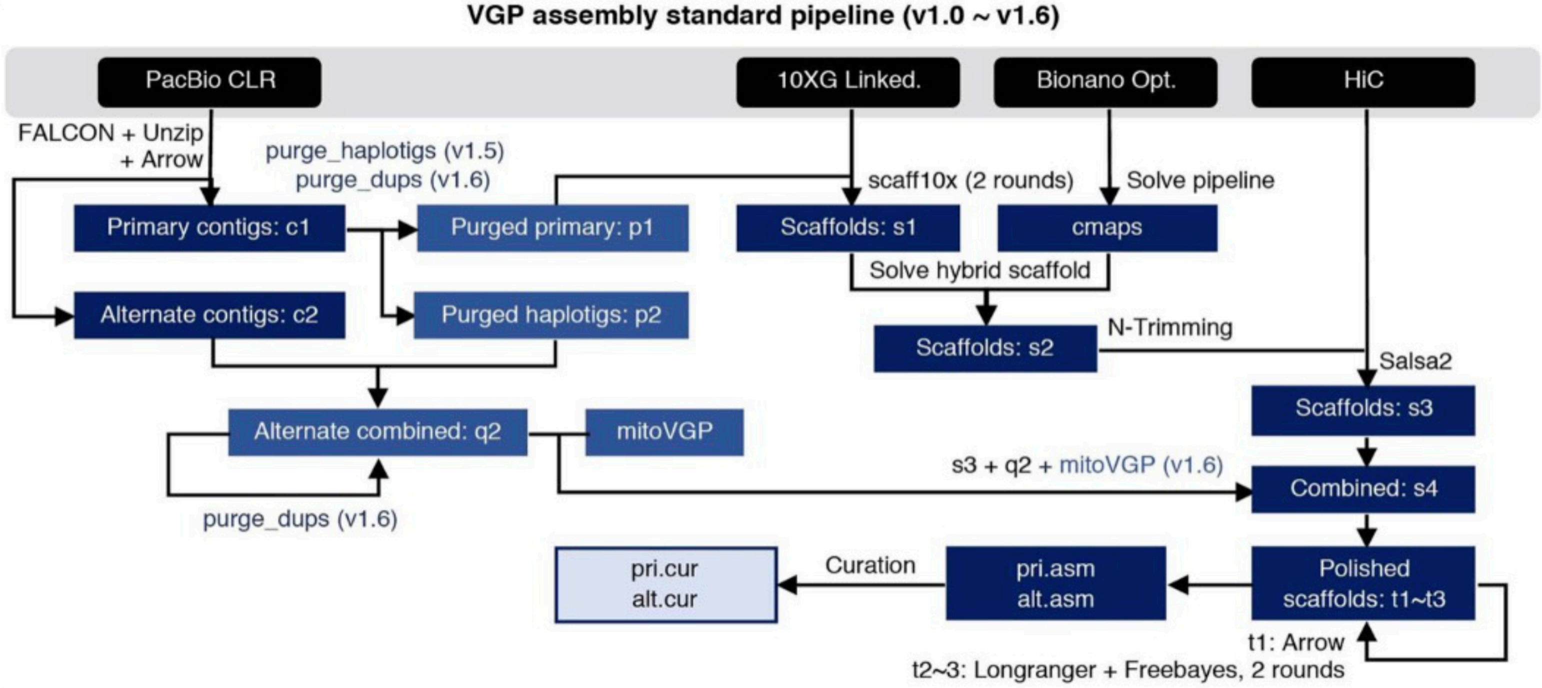


b

Taken from Rhie et al, 2021
