## Supplementary material for "Caecilian genomes reveal the molecular basis of adaptation, and convergent evolution of limblessness in snakes and caecilians": Figure 1

*Geotrypetes seraphini* Hi-C heatmaps before (left) and after (right) manual curation

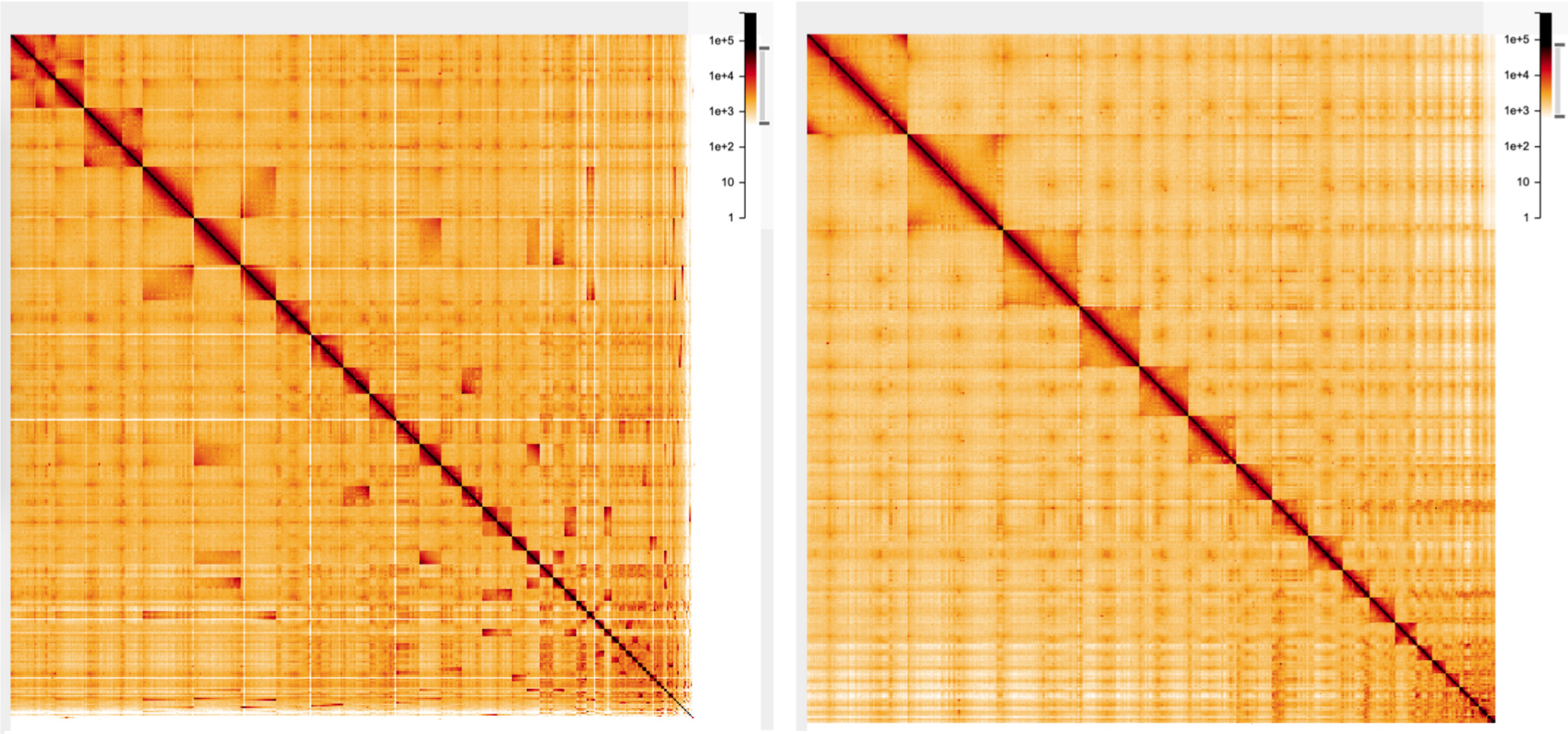


*Microcaecilia unicolor* Hi-C heatmaps before (left) and after (right) manual curation

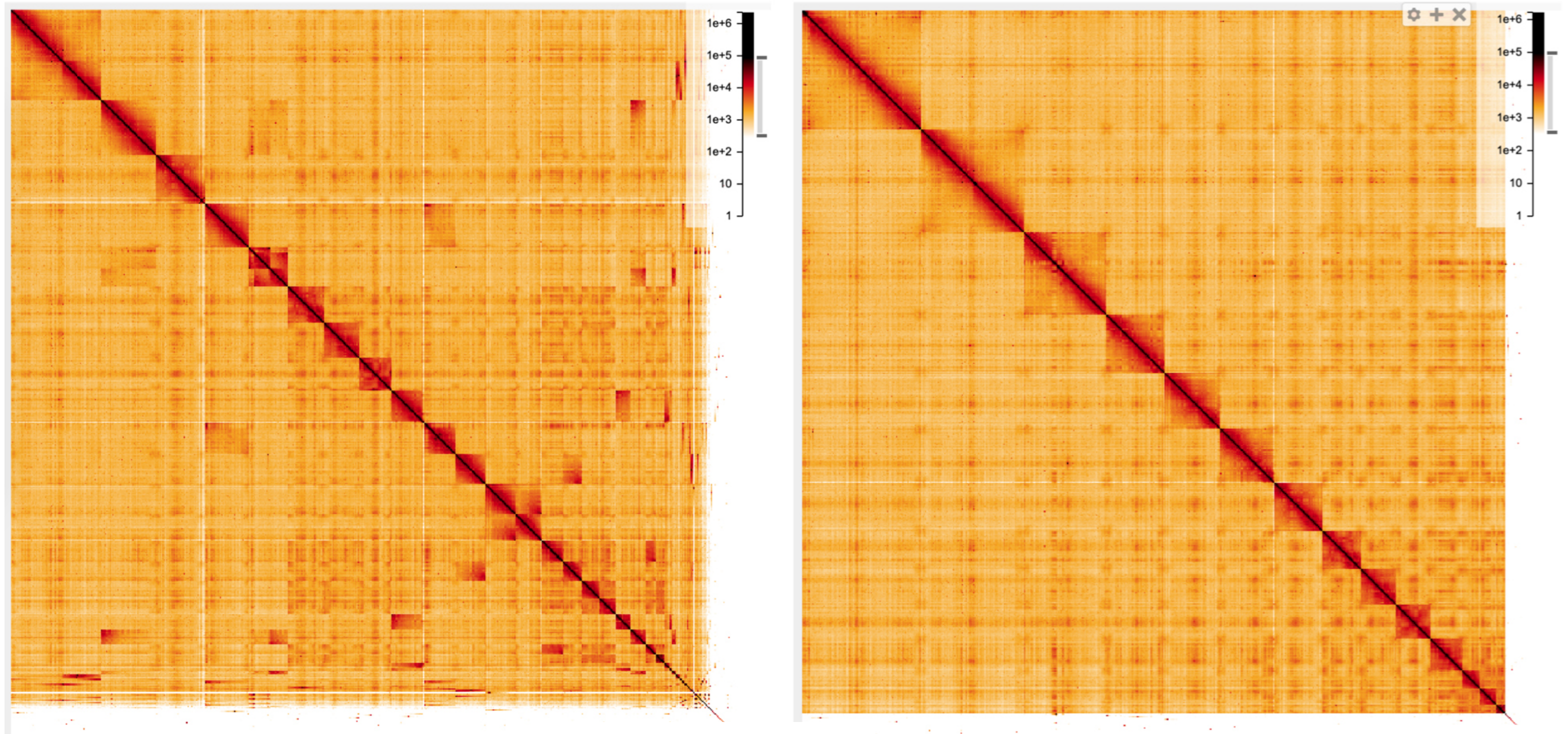
