## Supplementary Figure S2 for "Caecilian genomes reveal the molecular basis of adaptation, and convergent evolution of limblessness in snakes and caecilians"

Supplementary Figure S2: Alignment of ZRS enhancer region across a range of vertebrates illustrating the loss of an otherwise well conserved ZRS region in snakes and caecilians.

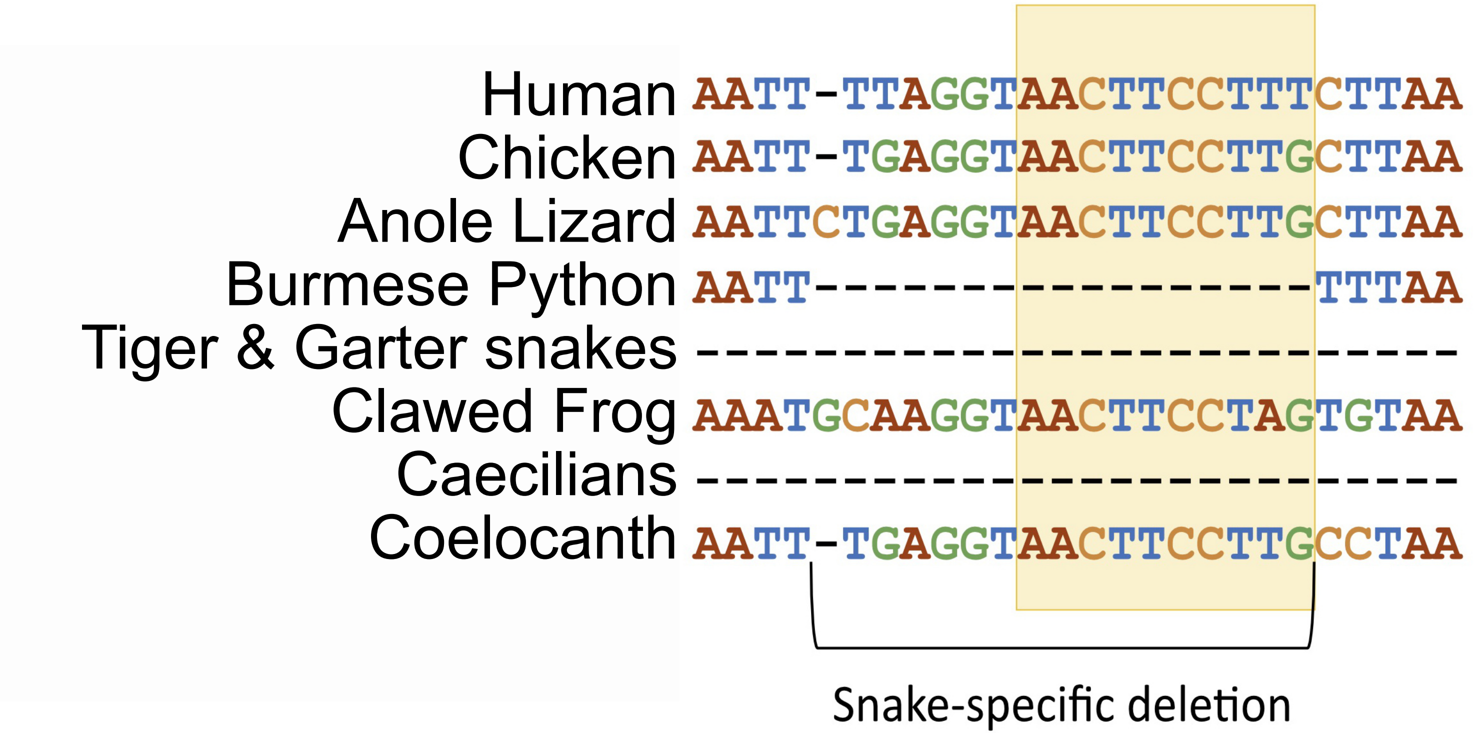
